# Strain-level analysis of *Bifidobacterium spp*. from gut microbiomes of adults of differing lactase persistence genotypes

**DOI:** 10.1101/2020.07.16.207811

**Authors:** Victor Schmidt, Hagay Enav, Timothy Spector, Nicholas D. Youngblut, Ruth Ley

## Abstract

One of the strongest associations between human genetics and the gut microbiome is a greater relative abundance of *Bifidobacterium* in adults with lactase gene *(LCT)* SNPs associated with lactase-non persistence (GG genotypes), versus lactase persistence (AA/AG genotypes). To gain a finer grained phylogenetic resolution of this association, we interrogated 1,680 16S rRNA libraries and 245 metagenomes from gut microbiomes of adults with varying lactase persistence genotypes. We further employed a novel genome-capture based enrichment of *Bifidobacterium* DNA from a subset of these metagenomes, including monozygotic (MZ) twin pairs, each sampled 2 or 3 times. *B. adolescentis* and *B. longum* were the most abundant *Bifidobacterium* species regardless of host *LCT*-genotype. *LCT-* genotypes could not be discriminated based on relative abundances of *Bifidobacterium* species or *Bifidobacterium* community structure. Metagenomic analysis of *Bifidobacterium-enriched* DNA revealed intra-individual temporal stability of *B. longum, B. adolescentis,* and *B. bifidum* strains against the background of a changeable microbiome. We also observed greater strain sharing within MZ twin pairs compared to unrelated individuals, and within GG compared to AA/AG individuals, but no effect of host *LCT*genotype on *Bifidobacterium* strain composition. Our results support a “rising tide lift all boats” model for the dominant Bifidobacteria in the adult gut: their higher abundance in lactase-non persistent compared to lactase-persistent individuals results from an expansion at the genus level. *Bifidobacterium* species are known to be transmitted from mother to child and stable within individuals in infancy and childhood: our results extend this stability into adulthood.

**IMPORTANCE:** When human populations domesticated animals to drink their milk they adapted genetically with the ability to digest milk into adulthood (lactase persistence). The gut microbiomes of lactase non-persistent people (LNP) differ from those of lactase-persistent people (LP) by containing more bacteria belonging to the Bifidobacteria. These beneficial gut bacteria, which fall into many species, are known to degrade milk in the baby gut. Here, we asked if adult LP and LNP microbiomes differ in the species of Bifidobacteria present. We studied the gut microbiomes of LP and LNP adults, including twins, sampled at several times. In particular, we used a technique to selectively pull out the DNA belonging to the Bifidobacteria: analysis of these DNA segments allowed us to compare Bifidobacteria at the strain level. Our results show that the LNP enhance the abundance of Bifidobacteria regardless of species. We also noted that a person’s specific strains are recoverable several years later, and twins tend to share the same ones. Given that Bifidobacteria are inherited from mother to child, strain stability over time in adulthood suggests long term, multi-generational inheritance.

## INTRODUCTION

Lactose tolerance arose in European, African and Middle Eastern human populations with animal domestication (Tishkoff *et al.*, 2007; Itan *et al.*, 2010; Ranciaro *et al.*, 2014). The genetic underpinnings of lactose tolerance represent one of the strongest signals of recent selection in the human genome. The enzyme lactase metabolizes lactose, the primary carbohydrate in mammalian milk. The gene regulatory region of the lactase gene (*LCT*) controls the downregulation of lactase after weaning (Sabeti *et al.*, 2007). Allelic variants that inhibit downregulation, resulting in lactase persistence, occur in an estimated 35% of humans (Itan *et al.*, 2010). Lactase persistence allows hydrolyzation of lactose and uptake of the resulting glucose and galactose directly in the small intestine of adults and is linked to lactose tolerance.

In a striking parallel, one of the strongest signals for human genotype effects on the gut microbiome also relates to lactase persistence. In Western populations, individuals with a lactase persistent genotype harbor significantly lower relative abundance of *Bifidobacterium* in their gut microbiomes compared to non-persistent individuals (Blekhman *et al.*, 2015; Goodrich *et al.*, 2016a; Turpin *et al.*, 2016; Rothschild *et al.*, 2018). The association was found to be stronger when dairy consumption in the lactase non-persistent individuals was considered (Bonder *et al.*, 2016). Together these observations suggest that for lactase non-persistent individuals, *Bifidobacterium* may benefit from the availability of lactose in the gut. *Bifidobacterium* is a large genus whose members partition the nutrient niche space (O’Callaghan *et al.*, 2016), which suggest that not all *Bifidobacterium* species may respond equally to lactose availability. However, beyond an overall enrichment of the *Bifidobacterium* genus, the effects of the lactase persistence genotype on the Bifidobacteria community in the gut remain unclear.

Here, we aimed to interrogate the *LCT-Bifidobacterium* link at a finer phylogenetic resolution. We re-examined public metagenomic (Xie *et al.*, 2016) and 16S rRNA gene data (Goodrich *et al.*, 2016a) from an UK twin cohort with both lactase persistent (AA, AG) and non-persistent (GG) individuals. We then performed genome-capture enrichment of *Bifidobacterium* from 11 twin pairs of each genotype across two or three time points per individual, and sequenced each metagenome before and after genome capture. With these data, we asked if lactase persistence genotype influenced strain composition, longitudinal stability within an individual, or similarity within families for three species (*B. longum, B. adolescentis, B. bifidum).* Our results suggest a proportional increase of the predominant *Bifidobacterium* species in the gut microbiomes of the lactase-persistent compared to non-persistent individuals. We observed strain sharing for twins, and stability within individuals, regardless of *LCT* genotype.

## RESULTS

Our analysis of 16S rRNA gene SVs confirmed our previous OTU-based report (Goodrich *et al.*, 2016a) of significantly greater mean relative abundance of *Bifidobacterium* in lactase non-persistent (GG) versus persistent (AA/AG) individuals (mean AA/AG = 0.96%±0.05 and mean GG = 3.22%±0.4SE, linear mixed-model *P* < 4.5e^-16^)(Figure 1A, Table S1). This analysis revealed 13 *Bifidobacterium* SVs which occurred in at least two of 1,680 samples. Five of the 13 SVs were unambiguously identified to the species level, while two fell within a broader *B. longum/B. breve* clade. Both host genotype groups (GG vs. AA/AG) were dominated, in order of abundance, by *B. adolescentis, B. longum/breve, B. pseudocantenulatum, B. animalis* and *B. bifidum.* Together these taxa represented over 98.6% of all sequences assigned to the *Bifidobacterium* genus, although only 1.1% of all sequences across all taxa (Table S1). These five species had similar proportions of the total *Bifidobacterium* community in both genotype groups and almost identical rank orders (GG: 93%, AA/AG: 91%) (Figure 1B, Table S1). Based on SVs, *B. animalis* and *B. dentium* were the only taxa with species level designations that did not have significantly greater relative abundance in GG versus AA/AG genotypes.

**Figure 1:**
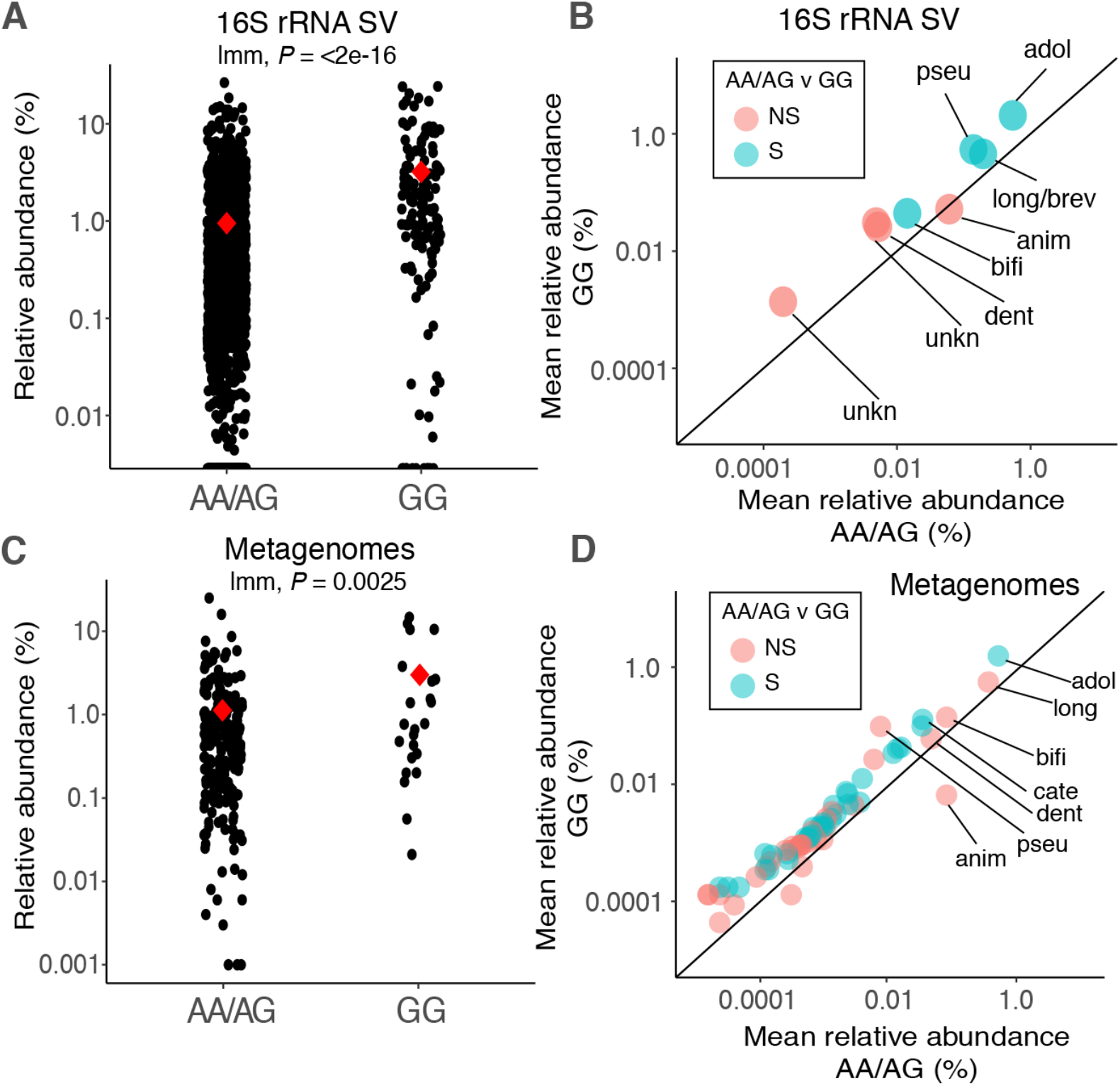
Lactase persistent genotype enriches most human associated *Bifidobacterium* species. A: Relative abundance of reads annotated as *Bifidobacterium* at the genus level from lactase persistent (AA/AG) and non-persistent (GG) individuals based on 16S rRNA SV taxonomic annotations (N: AA/AG = 1,549, GG = 131).B: Mean relative abundance of *Bifidobacterium* species in AA/AG (x-axis) and GG (y-axis) individuals based on 16S rRNA SV taxonomic annotations. Colors indicate significant enrichment in GG. 1:1 line designates equal proportion in each genotype, points above the line therefore indicate enrichment in GG and *vice versa.* Taxa that only occurred in one genotype are not shown. Species are; *B. adolescentis* (adol), *B. longum* (long), *B. pseudocatenulatum* (pseu), *B. animalis* (anim), *B. bifidum* (bifi), *B. dentium* (dent), *B. catenulatum* (cate), and unknown (unkn) indicates SVs that cannot be resolved beyond genus level. C: Relative abundance of reads annotated as *Bifidobacterium* at the genus level from lactase persistent (AA/AG) and non-persistent (GG) individuals metagenomic annotations (N: AA/AG = 222, GG = 23). Red diamonds indicate means and *P* values from linear mixed models. D: Mean relative abundance of *Bifidobacterium* species in AA/AG (x-axis) and GG (y-axis) individuals as revealed by metagenomic annotations. Colors as in B.

After normalization of *Bifidobacterium* within each genotype (thereby removing the effect of an overall enrichment of the genus), discriminant analysis using LEfSe (Segata *et al.*, 2011) revealed no discriminant *Bifidobacterium* SVs between GG and AA /AG individuals (data not shown). This result indicates similar proportional abundances of each *Bifidobacterium* species within each host genotype group, despite an overall greater relative abundance of all taxa in GG versus AA/AG genotypes. ANOSIM permutation tests on the Bray-Curtis Dissimilarity (BCD) matrix of within-genotype normalized *Bifidobacterium* SV matrixes also revealed no significant community clustering by host genotype group (ANOSIM *P* > 0.05), further supporting a proportional increase across most taxa rather than genotypic selection of specific species or strains within the genus.

We also assessed the association between frequency of dairy consumption and *Bifidobacterium* SVs for the 783 samples for which both datasets were accessible. Linear mixed-effect models, with genotype as a random variable, revealed no overall association between dairy consumption and the relative abundance of the genus (linear mixed model *P =* 0.113). When SVs were interrogated independently, *B. animalis* and an unclassifiable Bifidobacteria were the only two SVs which showed significant associations (*B. animalis: P =* 0.0023, *B. unknown1: P =* 0.014). Interestingly, when each genotype was considered independently, significant associations with levels of dairy consumption were observed for *B. animalis* within GG individuals, but not in AA/AG individuals (generalized linear model, GG: *P =* 0.03, AA/AG: *P =* 0.13).

Our taxonomic annotations of metagenomic reads from existing TwinsUK metagenomes (Xie *et al.*, 2016) revealed an overall enrichment of the *Bifidobacterium* genus in lactase-non-persistent individuals (mean AA/AG = 1.1%±0.15 and mean GG = 2.8%±0.9, linear mixed model *P =* 0.0025)(Figure 1C, Table S1) and a proportional increase across most *Bifidobacterium* species in GG versus AA/AG individuals (Figure 1D). Largely concordant with the results of the 16S rRNA SV analysis, Bifidobacteria metagenome annotations from both host genotype groups were dominated by *B. adolescentis, B. longum, B bifidum,* and *B. animalis,* which together represented more than 80% of all *Bifidobacterium* sequences, though only 1% of the overall community (Figure 1D, Table S1). Taxonomic annotations from metagenomes showed a greater diversity of *Bifidobacterium* taxa across the full dataset compared to the 16S rRNA gene SV based analysis, despite a lower sample count (245 metagenomes versus 1,680 16S rRNA samples), with 65 species and subdivisions identified (versus 13 SVs).

Functional annotations of the 245 metagenomes revealed no MetaCyc *Bifidobacterium* metabolic pathways with significantly different relative abundances between the two genotype groups (pairwise Kruskal-Wallis H-tests for mean abundance of each *Bifidobacterium* pathway in GG versus AA/AG individuals, Bonferroni multiple comparison correction, implemented in HUMAnN2; data not shown). However, low or zero read coverage for most *Bifidobacterium* pathways in the metagenomes limited the power to assess differences between host genotype groups at the functional level.

### Genome capture enriches Bifidobacteria DNA

We used genome capture (Carpi *et al.*, 2015; Metsky *et al.*, 2019) to enrich for *Bifidobacterium* DNA in metagenome libraries generated from 22 individuals each with 2 or 3 samples obtained at different time points (46 samples total representing 11 GG and 11 AA individuals). Each genotype group also included 4 sets of adult monozygotic twin siblings (Table S2). Our custom genome capture array included a total capture space of >94Mb and 89k capture targets built from 47 reference *Bifidobacterium* genomes spanning the entire diversity of the genus (Table S3). The capture reaction was largely genus-specific, with low levels of enrichment of *non-Bifidobacterium* Bifidobacteriaceae and other *non-Bifidobacterium* Actinobacteria (Figure 2A). The capture reaction increased the mean relative abundance of all *Bifidobacterium* across our metagenomic sample subset from 2.1% (±0.27%SE) to 60.2%(±3.7%SE), representing a nearly 30 fold increase from pre-to postcapture libraries. The mean relative abundance of sequences annotated as a specific *Bifidobacterium* taxa across all pre-captured libraries was proportional to that taxon’s initial relative abundance in the pre-captured libraries *(rho* = 0.99, *P* < 2.2e^-16^, Figure 2B), and the mean relative abundance ranking order of the top 5 taxa was the same in pre- and post-captured libraries (data not shown).

**Figure 2:**
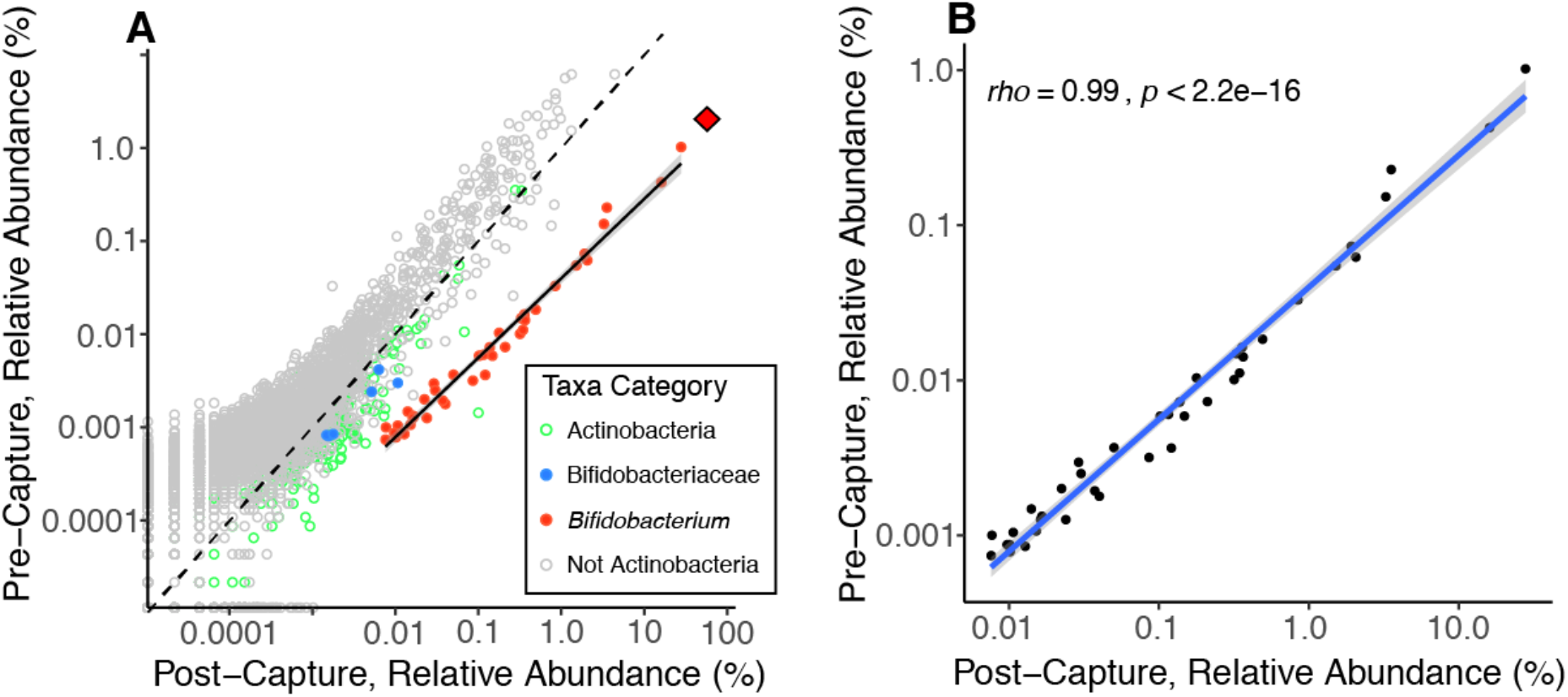
*Bifidobacterium* is proportionally enriched using genome capture. **A:** Relative abundance of all species level annotations from the genome capture subset (N = 46), with each point representing the mean relative abundance of a distinct species level classification before (pre-capture) and after the *Bifidobacterium-DNA* capture assay (post-capture). Points are colored by taxonomic relatedness to *Bifidobacterium.* Pre and post capture libraries from a single sample were shotgun-sequenced independently. The dotted 1:1 line represents equal relative abundances in pre and post capture libraries, points below the 1:1 line are enriched in post-capture libraries relative to pre-capture libraries. The solid black line shows the least squares regression between pre and post capture for the *Bifidobacterium* annotations. The shallower slope of the *Bifidobacterium* enrichment line versus the 1:1 line indicates greater enrichment of more common taxa. The red diamond shows the total relative abundance of *Bifidobacterium (i.e.* the sum of all *Bifidobacterium* annotations) in each of pre and post capture libraries. **B:** Same data as plot A but only showing *Bifidobacterium* annotations. The blue line shows least-squares regression with 95% confidence intervals, spearman’s rho is shown along with the significance of the correlation.

### Strain level analysis of *Bifidobacterium spp*. in post-capture metagenomes

We compared strains recovered from post-capture metagenomes using three methods: (i) we generated genome-wide strain phylogenies using multi-locus sequence alignments (MLSAs) via StrainPhlAn (Truong *et al.*, 2017); (ii) a marker-SNP strain tracking analysis implemented in the Metagenomic Intra-Species Diversity Analysis System (MIDAS)(Nayfach *et al.*, 2016); and, (ii) we used a custom synteny-based approach. We compared strains in all possible pairwise sample comparisons from our genome capture subset (N = 46). We then grouped each comparison into one of three categories; *‘Intra-Individual (comparison between samples from the same person over time), ‘Same Family’* (between monozygotic twins within a twinship), and *‘Unrelated’* (between samples from unrelated individuals). These categories allowed us to assess strain stability through time within an individual (intra-individual comparisons) and the influence of monozygotic twins (same family comparisons). Our post-capture libraries yielded sufficient reads to conduct these analyses in four species, *B. longum, B. adolescentis, B. animalis, and B. bifidum,* with sample numbers shown in Table 1.

**Table 1:** Sample number and comparison category overview for StrainPhlAn and MIDAS strain-level analyses.

|  | StrainPhlAn |  |  |  | MIDAS |  |  |  |
| --- | --- | --- | --- | --- | --- | --- | --- | --- |
|  | Samples with sufficient coverage for phylogeny | Number of 'intra-individual', 'Same Family', and 'Unrelated' comparisons |  |  | Samples with sufficient coverage for marker-SNP analysis | Number of 'Intra-individual', 'Same Family', and 'Unrelated' comparisons |  |  |
| *B. adolescentis* | 43 | 23 | 37 | 843 | 37 | 17 | 30 | 619 |
| *B. longum* | 39 | 20 | 29 | 692 | 36 | 15 | 24 | 591 |
| *B. bifidum* | 19 | 8 | 8 | 155 | 20 | 6 | 8 | 176 |
| *B. animalis* | 11 | 3 | 2 | 50 | 15 | 4 | 3 | 98 |
| **Total** |  | **54** | **76** | **1740** |  | **42** | **65** | **1484** |

### MLSAs showed greater similarities of strain sequences in intra-individual versus unrelated comparisons

Using StrainPhlAn, we created MLSAs across nearly 200 strain-resolving marker genes derived from a species’ core genome (genes shared by all strains in the species). Of the post-capture shotgun metagenomes, *B. adolescentis, B. longum, B. bifidum* and *B. animalis* had sufficient coverage for 43, 39, 19, and 11 samples respectively (minimum of 2x coverage across entire alignment) (Table 1). The marker gene MLSAs ranged from 36 and 83kb in length depending on species (Table S4). The mean number of polymorphic sites within a sample across an alignment ranged from 0.39% to 0.92%, while the dominant allele at each site ranged upwards from 76%, suggesting low strain diversity within samples (Table S4, Figure S1).

Mean patristic distance *(i.e.,* branch length) was significantly less in intra-individual versus unrelated comparisons in phylogenies of *B. adolescentis, B. longum,* and *B. bifidum* strains (bootstrapped Wilcoxon rank sum *P* < 0.05 in all three cases, Figure 3). This result indicates strain stability: an individual’s strains are more similar for the same person sampled over time than compared to strains from a different person. We did observe some instances of large patristic distance between samples of the same individual (Figure 4), and in both cases where three samples per individual were included, one of the three samples had a large patristic distance more typical of unrelated comparisons (Figure 4). These large inter-individual patristic distances could result from detection errors *(i.e.,* a strain was present in both samples but failed to be detected in one), or from differences in the relative abundance of strains differed between time points (this analysis picks the most abundant), or from loss and regain of strains.

**Figure 3:**
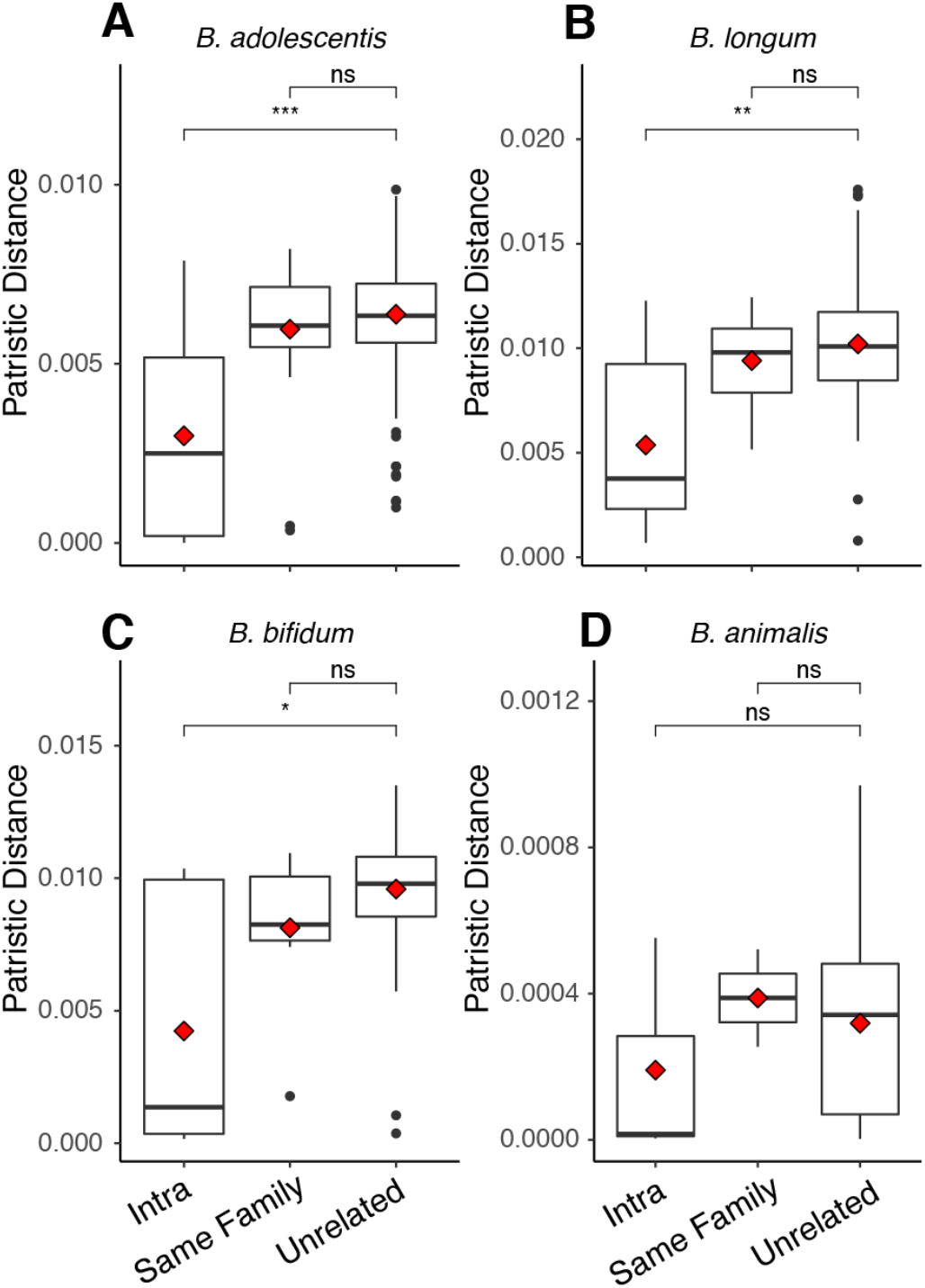
Patristic distances between samples from RAxML phylogenies of four strains shows intraindividual strain similarity. Patristic distances (branch lengths) between all samples within our StainPhlan phylogenies, with each point representing the distance between two samples on a single phylogeny. Distances are separated by the category of comparison; Intra (same person over time), Same Family (comparisons between twin siblings), and Unrelated (comparisons between unrelated people). Species are; A) *B. adolescentis,* B) *B. longum,* C) *B. bifidum,* and D) *B. animalis.* Boxplots show median and quartiles, while red diamonds show mean. Significance levels are median Wilcoxon rank sum tests after 999 bootstraps to smallest group size, ns: p > 0.05 *: p <= 0.05, **: p <= 0.01, ***: p <= 0.001, ****: p <= 0.0001.

Mean patristic distance of same-family comparisons was not significantly different than unrelated comparisons for any of the four taxa examined (Wilcoxon rank sum of mean distances in intertwin versus unrelated, *P* > 0.05 in all cases, Figure 3). Thus, the MLSA-based RaxML phylogenies did not show that co-twins harbored similar strains of these 4 species. Furthermore, this analysis did not reveal any significant grouping of strains according to lactase genotype (ANOSIM permutations of phylogenetic distance matrix *P >* 0.05, Figure 4) and longitudinal intra-individual comparisons were not significantly different between genotype groups for any of the four species we examined either individually (Figure S2) or in aggregate (Figure S3)(Wilcoxon rank sum test of mean intra-individual distance, AA versus GG, *P* > 0.05; Table 2A & S5). The time interval between repeated samples from the same individual and patristic distance did not reveal any relationship for any species (Spearman’s rho non-significant in all cases, Figure S4), supporting strain stability with time.

**Figure 4:**
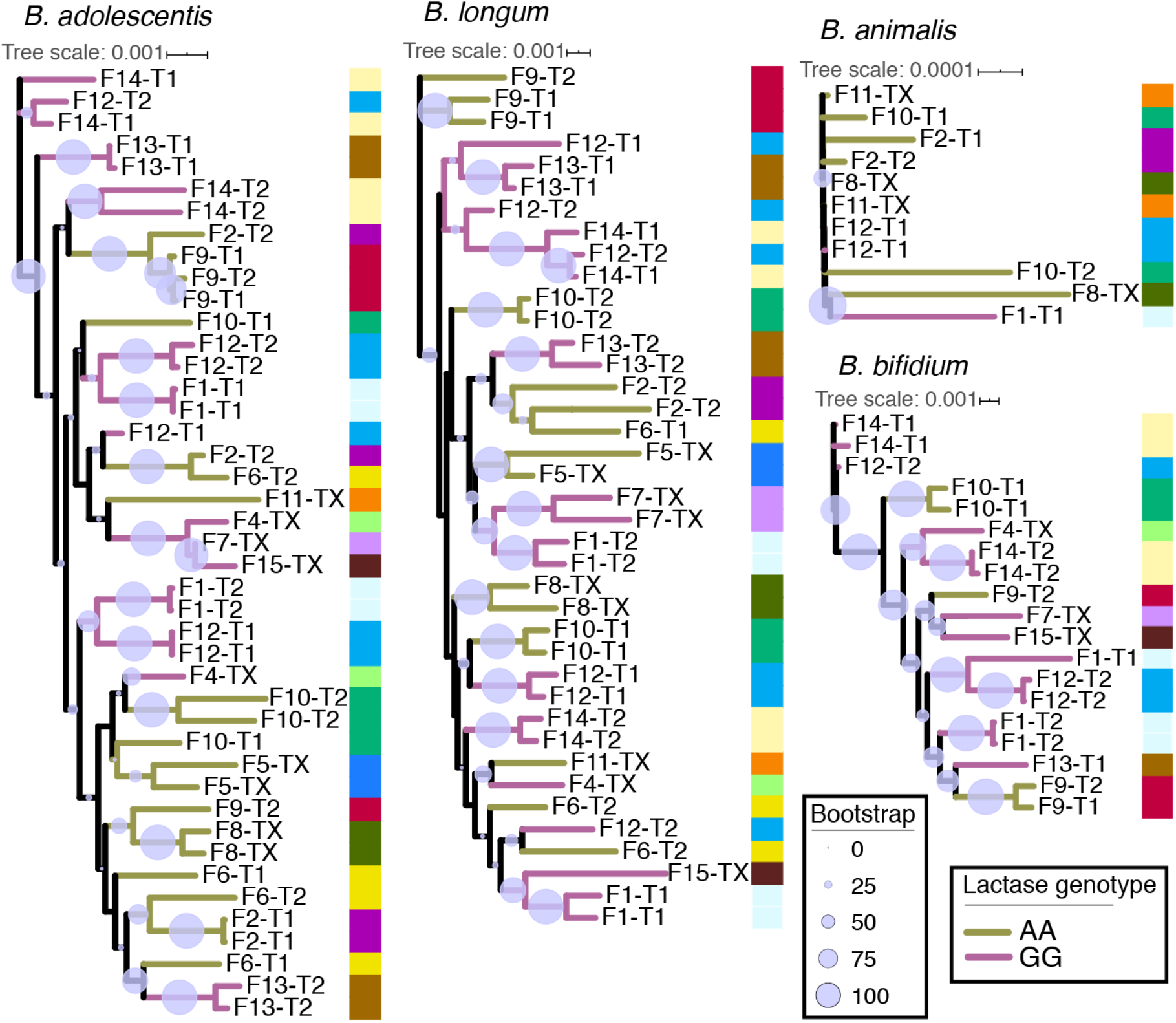
Strain phylogenies show longitudinal intra-individual strain similarities. RAxML phylogenies of StrainPhlAn MLSAs for each of the four species considered. Tree branches are colored by lactase persistence genotype. Columns of colors at each node reference family IDs, with multiple samples from the same individual and their twin having the same color. Samples are labeled by family ID, then by twin ID within the family. Identical labels imply samples from different time points of the same individual (i.e. intra-individual sample). Twin IDs marked with ‘X’ imply only a single twin had representation on the tree, or that individual had no twin in the dataset. Scales show patristic distances.

### Marker-SNPs are shared at higher percentages in intra-individual versus unrelated comparisons

To further interrogate *Bifidobacterium* strain stability within individuals over time, we employed a ‘strain tracking’ feature, implemented in the MIDAS software, to identify rare marker-SNPs highly discriminant for a specific strain in an individual at a single time point. We had sufficient coverage to employ this approach in >10 individuals for each of *B. longum, B. adolescentis, B. animalis,* and *B. bifidum* species (Table 1). Our results show that microbe marker-SNPs of an individual are far more likely to be found in the same individual at a different time point than in an unrelated individual in three of the four species examined (bootstrapped Wilcoxon rank sum of mean %shared SNPs, intra-individual versus unrelated, *P* < 0.0001 in all cases, Figure 5). *B. animalis* was the one exception, where SNP sharing was not significantly higher in intra-individual versus unrelated comparisons (bootstrapped Wilcoxon rank sum *P =* 0.32, Figure 5; Tables 2B & S5).

**Figure 5:**
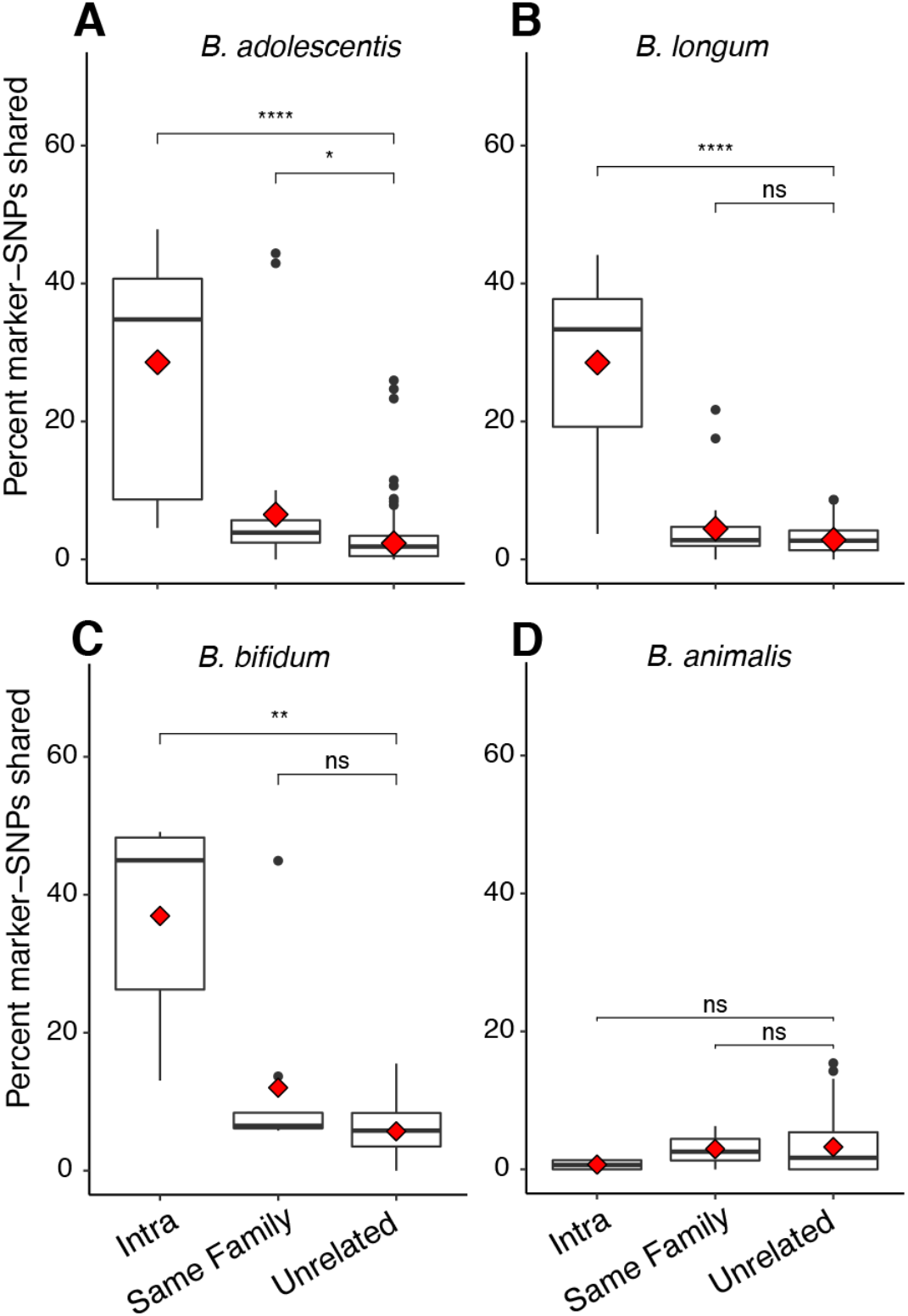
Marker-SNP sharing over time and between individuals shows strain stability and twin sharing in three of four species. Percent of marker-SNPs shared for all pairwise comparisons, separated by the category of comparison; Intra (same person over time), Same Family (comparisons between twin siblings), and Unrelated (comparisons between unrelated people). Shown are the four taxa for which sufficient coverage existed to calculate marker SNPs in at least six individuals, species are; A) *B. adolescentis,* B) *B. longum,* C) *B. bifidum,* and D) *B. animalis.* Boxplots show median and quartiles, while red diamonds show mean. Significance levels are median Wilcoxon rank sum tests after 999 bootstraps to smallest group size, ns: p > 0.05 *: p <= 0.05, **: p <= 0.01, ***: p <= 0.001, ****: p <= 0.0001.

We observed greater mean intra-individual sharing of *B. adolescentis* and *B. longum* in GG versus AA individuals (Figure S2), when all species were considered in aggregate (Wilcoxon rank sum of mean intra-individual %shared SNPs, GG versus AA individuals, *P =* 3.3e^-9^; Figure S3). This pattern was driven by *B. longum* and *B. adolescentis,* the two most abundant species (Figure S2).

The SNP-marker analysis revealed that high stability in either *B. longum, B. bifdum,* or *B. adolescentis* was a good predictor of stability in the other two species within that same individual (Figure 6). This suggests some individuals carry stable strains, while others witness strain replacement, across all three species together. We detected higher marker-SNP sharing within families *(i.e.,* a twin effect) for *B. adolescentis* (bootstrapped Wilcoxon rank sum *P =* 0.021) but not the other three species examined, possibly due to a lower power to detect this pattern. The percent of shared marker-SNPs for a given intra-individual comparison was not correlated with time between sample collection, nor where other metrics of overall microbiome similarity (Figure S5), meaning the percent of shared marker-SNPs between samples does not decrease with time, and the patterns observed here are therefore not a function of time-induced SNP accumulation.

**Figure 6:**
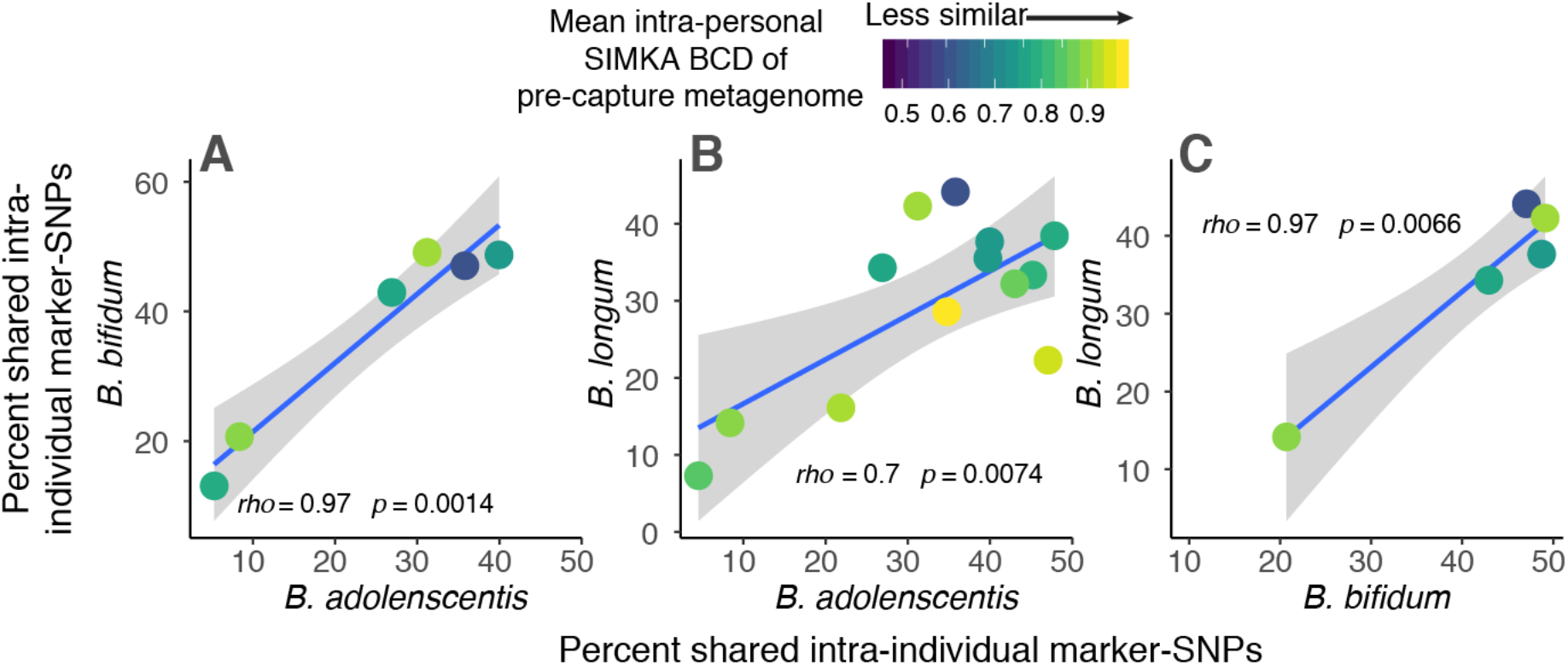
Intra-individual marker-SNP sharing across different *Bifidobacterium* species. Each plot shows the percent of marker-SNPs shared within an individual over time for two different *Bifidobacterium* species, each of three comparison possibilities are show across plots A, B, and C. Each point represents a comparison between longitudinal samples from the same individual. Points are colored by the Simka Brey-Curtis Dissimilarity (BCD) value between the two samples, as measured from pre-captured metagenomes, and is therefore a measure of overall microbiome dissimilarity over time within an individual. Blue line shows least-squares regression with 95% confidence intervals, and spearman’s rho along with the significance of the correlation is shown.

Notably, *Bifidobacterium* strains exhibited intra-individual longitudinal stability despite high variability in overall microbiome communities around them. Only a very weak, non-significant association was observed between an individual’s overall microbiome stability (assessed using Brey-Curtis Dissimilarity (BCD) of pre-captured metagenomes) and the number of shared marker SNPs for a given *Bifidobacterium* strain (Figure 6, Figure S6). This pattern held when microbiome community dissimilarity was assessed using either Simka BCD values, a k-mer based annotation-independent method, or BCD values for annotations of metabolic pathways of pre-captured metagenomes (Figure S6). These results suggest *Bifidobacterium* stability may be independent from the dynamics of microbiome communities in which they exist.

### Synteny analysis reveals sharing of strains within twin pairs and strain persistence in individuals over time

The analyses described above are based on single nucleotide variations and gene-content comparisons. Gene synteny is defined as the conservation of gene order between chromosomes, along an evolutionary gradient (Engström *et al.*, 2007) and was previously used as a measure of the distance between genomes (Nadeau and Taylor, 1984; Sankoff *et al.*, 1992). Here, we identify syntenic blocks, regions of conserved DNA sequence occurring in the same order in two different chromosomes, to determine the relatedness of *Bifidobacterium* strains. We created a pipeline to compare the synteny of different genomic regions in three *Bifidobacterium* species for which we had sufficient coverage from captured metagenomes: *B. longum, B. adolescentis* and *B. bifidum* (see methods). Briefly, we performed a *de-novo* assembly of post-captured metagenomes in each sample, and used a subset of randomly selected genes to perform a BLAST search against the metagenomic assemblies. Next, we performed a pairwise synteny comparison between all genomic regions, which included the gene used as BLAST query and their flanking regions (3.5 kbp upstream and downstream to the gene). Finally, we defined a synteny-score and performed the same pairwise comparisons as outlined above for regions which were identified in >15 single-individual assemblies. In summary, we analyzed 10, 7 and 4 regions for *B. longum, B. adolescentis* and *B. bifidum,* respectively.

For all three species, intra-individual comparisons yielded significantly higher synteny scores than comparisons of unrelated individuals (Figure 7), although this pattern varied somewhat by region; 6/7 genomic regions for *B. adolescentis,* 2/4 regions for *B. bifidum,* and 4/10 regions for *B. longum* (results are summarized in Table 2C and *P-* values are shown in Table S6). These results indicate that the same strains persist in the gut over time. Comparisons of synteny between individuals within a twin pair yielded significantly higher scores (relative to comparisons of unrelated individuals) in 4 regions in *B. adolescentis* and in one region in *B. bifidum.* In *B. longum,* although not statistically significant, 3 regions showed lower synteny scores for individuals within a pair (P-values 0.055-0.08, Table S6). Finally, *LCT*-genotype did not influence longitudinal strain stability as measured by synteny. No differences were seen in mean intra-individual synteny scores between lactase persistence genotypes (Wilcoxon rank sum test of mean intra-individual synteny, AA versus GG, *P* > 0.05), and each genotype independently showed significant differences between intra-individual and unrelated comparisons (bootstrapped Wilcoxon rank sum test, *P =* 0.0005 for AAs and *P* = 6.1e^-5^ for GGs, Figure S3). Together, these results support strain sharing events within the families and maintenance of strains over time within individuals.

**Figure 7:**
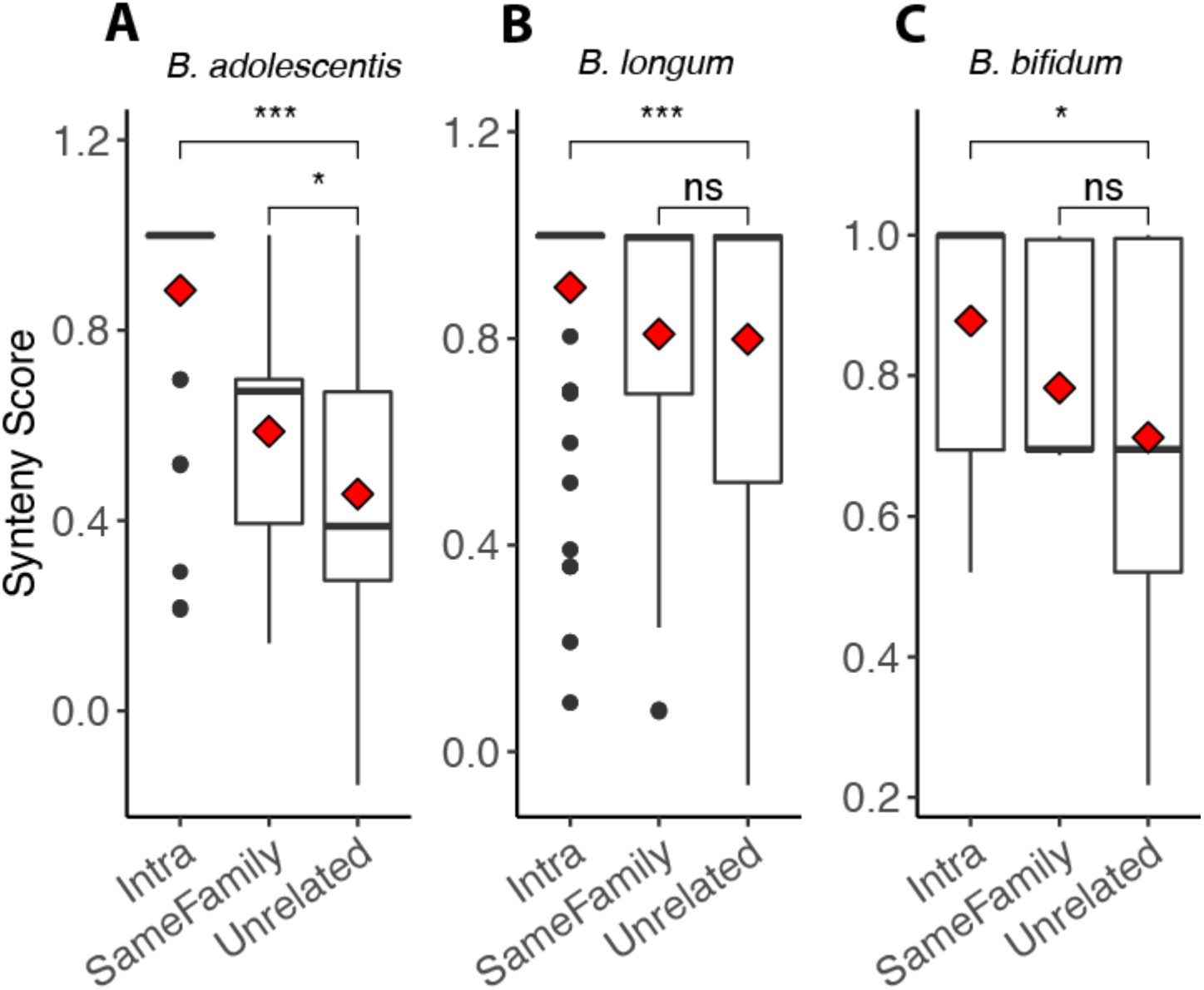
Within-species synteny scores over time and between individuals. Pairwise synteny scores for genomic regions of the three *Bifidobacterium* species for which sufficient read depth was generated for synteny analyses. Plots show all pairwise comparisons, separated by the category of comparison; Intra (same person over time), Same Family (comparisons between twin siblings), and Unrelated (comparisons between unrelated people). Species are; A) *B. adolescentis,* B) *B. longum,* C) *B. bifidum.* Boxplots show median and quartiles, while red diamond shows mean. For each species all regions were combined into a single analysis. Significance levels are Wilcoxon rank sum tests, ns: p > 0.05 *: p <= 0.05, **: p <= 0.01, ***: p <= 0.001, ****: p <= 0.0001.

**Table 2:** Overview of analyses for each of the four species examined at the strain-level, and for all species combined in aggregate. Panels A-C show significance for our three strain-level approaches. StrainPhlAn results are based on best tree values from RAxML hill-climbing algorithm. Within each panel, ‘Longitudinal stability’ refers to significantly greater similarity of Intra (same person over time) versus Unrelated (between unrelated people) comparison categories. ‘Stability GG > AA/AG’ refers to greater mean ‘Intra’ values for GG individuals versus mean ‘Intra’ values for AA/AG individuals *(i.e.,* is there greater stability within lactase non-persistent versus lactase persistent individuals). Finally, ‘Twin effect’ refers to greater similarity of Same Family (between twin siblings) versus Unrelated comparison categories. Panel D shows significance of genotypic enrichment from 16S rRNA SV data across the broad TwinsUK dataset (N = 1,680). In each panel ‘yes’ and ‘no’ refer to significance of the statistical test as described in the main text, while ‘NA’ indicates 3 or fewer samples were available in at least one category. In panel C (synteny) the number of significant regions out of the total number of regions with sufficient comparisons are shown. P-values are shown in Table S5 and S6. Species abbreviations are Adol: *B. adolescentis,* Long: *B. longum,* Bifi: *B. bifidum,* and Anim: *B. animalis.*

|  |  |  | adol | long | bifi | anim | In aggregate |
| --- | --- | --- | --- | --- | --- | --- | --- |
| **A** | Marker-gene based phylogeny (StrainPhlAn) | Longitudinal stability | yes | yes | yes | no | yes |
|  |  | Stability GG > Stability AA/AG | no | no | NA | NA | no |
|  |  | Twin effect | no | no | no | no | no |
| **B** | % of shared Marker-SNPs (MIDAS) | Longitudinal stability | yes | yes | yes | no | yes |
|  |  | Stability GG > Stability AA/AG | yes | yes | NA | NA | yes |
|  |  | Twin effect | yes | no | no | no | no |
| **C** | Synteny (all regions for each species in aggregate) | Number of gene regions | 7 | 10 | 4 | 0 | 21 |
|  |  | Longitudinal stability | 6/7 | 4/10 | 2/4 | NA | yes |
|  |  | Stability GG > Stability AA/AG | 0/2 | 0/8 | 0/0 | NA | no |
|  |  | Twin effect | 4/7 | 0/8 | 1/4 | NA | no |
| **D** | Enriched in GG versus AA/AG (16S rRNA SVs) |  | yes | yes | yes | no | yes |

## DISCUSSION

One of the strongest signals of host genetic effects on microbiome communities is the association between *Bifidobacterium* and a SNP in the regulatory region of the lactase gene *LCT*. Several independent studies have noted a greater relative abundance of the genus *Bifidobacterium* in the gut microbiomes of lactase non-persistent compared to lactase-persistent individuals of European origin (Blekhman *et al.*, 2015; Bonder *et al.*, 2016; Goodrich *et al.*, 2016a; Rothschild *et al.*, 2018). We did not detect an interaction between dairy consumption, *LCT* genotype and levels of *Bifidobacterium* here, as reported by Bonder (2016). However, our results show that compared to the lactase-persistent genotypes (AA/AG), the lactase non-persistence genotype (GG) enhances the proportion of prevalent and abundant *Bifidobacterium* species, without bias towards particular species, strains, or genome content. Our results also indicate that strains of certain *Bifidobacterium* species are shared within twin pairs and persist over time within individuals against a background of a dynamic microbiome community.

Corroborating results of previous reports (Turroni *et al.*, 2009; Arboleya *et al.*, 2016; Kato *et al.*, 2017), we observed that *B. adolescentis* and *B. longum* dominated the *Bifidobacterium* communities of the adult gut microbiome. Strain comparisons indicated that *B. adolescentis, B. longum* and *B. bifidum* were stable within adults over multi-year timescales and shared within twin pairs. It is interesting to note that *B. animalis,* the one species for which stability was not shown, was also among the few species not enriched in GG versus AA/AG genotype groups from our broader 16S rRNA and metagenomic surveys. These observations may reflect a different ecology of *B. animalis* compared to the others. *B. animalis* is commonly isolated from diary and other sources outside the human microbiome (Sun *et al.*, 2015), and has a reduced genetic repertoire for carbohydrate metabolism versus other taxa in the genus (Milani *et al.*, 2015b). *B. animalis* may therefore be a more transient autochthonous member of the adult gut microbiome.

A leading hypothesis for the greater relative abundance of *Bifidobacterium* in lactase non-persistent individuals is a greater availability of undigested lactose in the large intestine, which may be preferentially used by Bifidobacteria (Bonder *et al.*, 2016; Goodrich *et al.*, 2016b). The availability of undigested lactose in the guts of lactase non-persistent individuals could in principle result in niche partitioning between the different species. Indeed, many taxa within the *Bifidobacterium* genus are considered specialists (Turroni *et al.*, 2009; Milani *et al.*, 2015), and some devote significant proportions of their genomes to particular host derived resources including human milk oligosaccharides in *B. longum* subs. *infantis* (Sela *et al.*, 2008), and mucin glycans in *B. bifidum* (Duranti *et al.*, 2015). *B. longum* subs. *Infantis,* for example, is known to specialize in human milk oligosaccharide metabolism, while *B. longum* subs. *longum* is specialized in metabolism of plant-derived carbohydrates (Sela *et al.*, 2010; O’Callaghan and van Sinderen, 2016). Furthermore, gut *Bifidobacterium* taxa are known to follow successional patterns with a gradual shift from *B. longum* subs. *infantis, B. breve,* and *B. bifidum* in infant guts to *B. catenulatum, B. adolescentis* and *B. longum* subs. *longum* in adults (Arboleya *et al.*, 2016; Kato *et al.*, 2017). These shifts are thought to result in part from shifts in diet, and are associated with breastfeeding versus formula, weaning, and intake of solid food (Stewart *et al.*, 2018; Vatanen *et al.*, 2019). However, our results instead point to lactose utilization as enhancing all the common *Bifidobacterium* species, without altering their population structure. This implies that in the adult gut microbiome, common *Bifidobacterium* species use lactose without competing for it and it does not become a limiting resource. The non-differentiating effect of host *LCT*-genotype on the *Bifidobacterium* community results suggest a ‘rising tide raises all boats’ scenario, where a lactose utilization advantage is equally distributed among common *Bifidobacterium* species.

The strain comparisons in this study relied on enrichment of *Bifidobacterium* DNA by genome capture. This worked well for the most abundant species, *B. adolescentis* and *B. longum,* and allowed a reduced set of comparisons possible for *B. bifidum* and *B. animalis* due to lower coverage. To compare strains of the other Bifidobacteria present would require deeper sequencing or a more tailored set of probes in the genome capture. To compare strains we used two published methods based on sequence composition; one using an alignment of marker-genes (StrainPhlAn), and another using individually discriminant marker-SNPs (MIDAS). The novel synteny-based method that we developed allows for the identification of differences in the organization of genomic regions even when the sequence similarity is high. Generally, the two published methods agreed with the synteny approach, with some discrepancies (Table 2). All three methods detected intra-individual longitudinal stability for *B. adolescentis, B.bifidum andB. longum,* while no method detected stability in *B. animalis.* The three methods were divided however in their detection of within-family sharing of strains (a twin effect), and on the influence of lactase persistence genotype on strain stability. Both the marker-SNP and synteny approaches detected greater similarity of *B. adolescentis* strains between twins versus unrelated individuals, while our marker gene alignments did not. Only the synteny approach detected twin sharing in *B. longum* and *B. bifidum.* The marker-SNP approach was the only method to reveal any influence of lactase persistence on longitudinal stability, and did so only in *B. adolescentis* and *B. longum.* Why each method varied in its ability to detect a particular pattern is not clear, however their agreement with regards to longitudinal stability of *B. adolescentis, B. longum* and *B. bifidum* gives strong support to the conclusion that these species are stable within adults over multi-year timescales.

Temporal stability, along with vertical transmission within families, are expected to be associated with heritability. In addition to temporal stability and within-family sharing of strains show here, we had previously noted that the genus *Bifidobacterium* was both heritable and stable over time, based on 16S rRNA gene analysis (Goodrich et al., 2014). For the relative abundances of microbial taxa or genes to be stably heritable over time, they must be present every generation and associated with host genetic variation. If the acquisition of microbes from the environment or other individuals is less reliable than acquisition from parents, the heritability of horizontally acquired microbes should be less stable over time than the heritability of vertically acquired microbes. These results, together with previous observations, continue to reveal the Bifidobacteria as highly human-adapted members of the microbiome.

*B. adolescentis, B. longum* and *B. bifidum* have also previously been shown to be temporally stable in the gut microbiomes of infants, where they are far more abundant than in the adult gut microbiome. Strains of these species were stable within children for up to three years of age across diverse geographic cohorts (Vatanen *et al.*, 2019), and *B. longum* supbsp. *longum* was stable from infancy until six years of age (Oki *et al.*, 2018). Furthermore *B. bifidum* and and *B. longum* are transmitted from mother to infant (Asnicar *et al.*, 2017)(Yassour *et al.*, 2018). By extending existing evidence of *Bifidobacterium* strain stability to an adult cohort and demonstrating strain sharing within families, our results contribute to an overall picture of this genus, one which suggests long term maintenance of specific strains within an individual which can be transmitted across generations.

## METHODS

### Sample inclusion and access

All existing samples from the TwinsUK cohort were included assuming they had both 1) metagenomic or 16S rRNA sequence data and 2) genotype data at the 13910 G/A allele. All work involving the use of these previously collected samples was approved by the Cornell University IRB (Protocol ID 1108002388).

### 16S rRNA gene based community analyses

16S rRNA reads were extracted from existing datasets (N = 1,680) and analyzed via the Qiime2 pipeline (Bolyen *et al.*, 2019) with minor deviations. Briefly, PCR amplicons for the V4 region of the 16S rRNA gene were generated with primers 515F–806R and were sequenced with the Illumina MiSeq 2 × 250 v2 Kit at the Cornell University Institute for Biotechnology as previously described (Goodrich *et al.*, 2016a). DADA2 (Callahan *et al.*, 2016) was used to call 100% sequence identity Sequence Variants (SVs, a.k.a. 100% OTUs or ASVs). Taxonomy was assigned to SVs with the QIIME2 q2-feature-classifier (Bokulich *et al.*, 2018) using the SILVA database (v119)(Quast *et al.*, 2013). Taxonomic annotations of 16S rRNA Sequence Variants (SVs) were improved from the standard DADA2 output by extracting each SV representative sequence and querying it against the NCBI type strain database. In one case all NCBI annotations fell within the *B. longum/B. breve* clade, and was thus assigned as *‘B. longum/breve’* in subsequent analyses; in all other cases the original DADA2 assignment was kept.

Statistical testing for the influence of genotype was done using a non-parametric Wilcoxon ranksum test of the total relative abundance of all *Bifidobacterium* SVs between genotypes. Only SVs that occurred in at least 2 individuals across the dataset of 1,680 samples were included. Individuals with either AA or AG at 13910*A (rs4988235) were considered lactase persistent, while only GG was considered lactase non-persistent. Independent Wilcoxon rank sum tests were run for each *Bifidobacterium* SV independently across the two genotype groups. Because our dataset included replicate samples from the same individual (therefore not independent), we determined the overall influence of genotype on *Bifidobacterium* SV relative abundance using a linear mixed-effect model, with participant ID as a random effect, in the R package “lmr4”: *BifidobacteriumRA* ~ Genotype + (1 | ParticipantID).

To assess the community structure of the genus without the influence of an overall enrichment in GG individuals, all SVs annotated as ‘bifidobacterium’ were extracted into new SV tables and again normalized by 1 within an individual, then input into LEfSE (Segata *et al.*, 2011) via the Galaxy web interface with default parameters, genotype set as class, and no subclass. Longitudinal analyses of *Bifidobacterium* SVs narrowed the total sample count to 556 (278 matched pairs), and Spearman’s rho was calculated assigning time 1 and time 2 to one sample or another at random. To overcome unequal sample sized (GG = 30, AA/AG = 248), a mean Spearman’s rho was calculated after subsampling to N = 15 for each genotype over 999 permutations. All analyses were done in RStudio (v. 1.0.136).

Dairy intake was assessed as the portion size frequency for dairy for a week, adjusted for energy intake, using food frequency questionnaires developed and described elsewhere (Spector *et al.*, 2006). Associations with *Bifidobacterium* SVs were conducted using linear mixed-effect models, with genotype as a random effect, in the R package “lmr4”: *BifidobacteriumRA* ~ DairyFrequency + (1 | ParticipantID). Statistical significance was assigned using Satterthwaite’s method in the R package “lmerTest” (Kuznetsova *et al.*, 2017).

### Metagenomic community analyses

Metagenome sequences were used from existing datasets (N = 245) which were extracted as above with library preparation previously described (Karasov *et al.*, 2018). Taxonomic assignments were made using Kraken2 (Wood and Salzberg, 2014). We assessed the influence of genotype on the relative abundance of *Bifidobacterium* annotations using a mixed effect model as described above. Metabolic pathway annotations were generated using the HUMAnN2 pipeline against the MetaCyc database (Franzosa *et al.*, 2018), and significantly discriminant gene pathways were revealed using HUMAnN2’s built-in ‘humann2_associate’ script with default parameters at the gene pathway level (Franzosa *et al.*, 2018).

### Subset for genome capture, strain level analyses, and discriminant functional pathways

An additional subset of the TwinsUK cohort was selected for genome capture based on the i) availability of genotype data at the lactase persistence gene (rs4988235) and ii) at least 2 longitudinal samples between 8 months and 4 years apart with a BMI change of less than 3. We note that variation in temporal distance between sampling events had no impact on stability, community composition, or synteny values (Figure S4, Figure S5). In total 20 individuals with 2 time points and 2 individuals with 3 time points were selected (a total of 46 samples across 11 GG individuals/ 11 AA individuals). Each genotype also contained 4 sets of twin siblings (Table S2). DNA samples were brought through metagenome creation, capture reaction, and sequencing according to NimbleGen (Madison, WI, USA) SeqCap EZ HyperCap Workflow v.1.0. Briefly, gDNA underwent enzymatic fragmentation and adapter mediated PCR using KAPA HyperPlus Library Preparation, followed by a 16h hybridization with a custom set of biotinylated long oligonucleotide probes (the ‘probe array’), followed by a final re-amplification. The probe array was designed and manufactured by NimbleGen and included overlapping coverage of a total capture space of >94Mb and 89k capture targets which covered 47 type strain *Bifidobacterium* genomes (Table S3). Pre-captured and post-captured metagenomes were then sequenced across two Illumina paired-end 300 cycle (HiSeq 3000). Sequences of pre and post captured metagenomes are available on the European Nucleotide Archive accession number PRJEB38000.

Analyses on each pre- and post-captured library from our 46 sample longitudinal subset (92 metagenomes total) were conducted with Simka, an annotation independent k-mer based metric (Benoit *et al.*, 2016), and annotation based HUMAnN2 pipeline against the MetaCyc database (Franzosa *et al.*, 2018). Simka Bray Curtis Dissimilarity (BCD) matrixes were calculated directly within the software. HUMAnN2 BCD matrixes were generated from gene pathway relative abundance tables, after removal of collapsed pathway stratifications, using vegdist in the Vegan R package (V. 1.0.136). Each pairwise comparison within the BCD matrixes was then classified into one of three categories: Intra-individual (same person over time), Same Family (comparisons between twin siblings), and Unrelated (comparisons between unrelated people). Wilcoxon rank sum tests were used to determine differences between comparison categories. To overcome issues of non-independence related to multiple samples from the same individual within the dataset, we ran 9,999 permutations of Wilcoxon Rank Sum tests using subsets of individuals equal to the smallest number of samples from either category, and reported the mean value. Finally, ANOSIM tests of BCD hierarchical clustering was conducted using the ‘anosim’ script part of the ‘vegan’ R package with 9,999 permutations. ANOSIM permutation tests were run on intra-individual clusters, unrelated clusters, and same family clusters.

### MIDAS Marker-SNPs and StrainPhlAn MLSAs

Strain level assessments were made using the Metagenomic Intra-Species Diversity Analysis System (MIDAS)(Nayfach *et al.*, 2016) and StrainPhlAn (Truong *et al.*, 2017). MIDAS’s ‘strain_straintracker.py’ program was used with default settings to identify marker-SNPs as previously described (Nayfach *et al.*, 2016). Briefly, MIDAS initially identifies abundant species by mapping unassembled metagenomic reads to a database of 30,000 reference genomes and then identifies per-species SNPs for each abundant species within a sample. To identify the SNPs used in our analyses, we had MIDAS identify rare, sample-discriminatory SNPs from a pool of 1 sample per individual *(i.e.* SNPs that were unique to that individual). We term these sample-discriminatory SNPs as ‘marker-SNPs’. Each individual’s additional sample(s) was then added back into the pool, and marker-SNP overlap was assessed between sets of two samples in all possible pairwise comparisons across the dataset. The percent of shared marker-SNPs was the number of marker-SNPs shared between two samples, divided by the total number across both individuals. Bootstrapped Wilcoxon rank sum tests were performed on the mean percent of marker SNPs shared in each comparisons category described above *(i.e.* Intra-individual, Same Family, and Unrelated). Bootstrapping was done down to the lowest number in a single category for any given comparison and run for 9,999 permutations, and the mean p-value was reported.

StrainPhlAn multi-locus sequence alignment (MLSA) RaxML phylogenies were created using default StrainPhlAn parameters as described elsewhere (Asnicar *et al.*, 2017; Truong *et al.*, 2017). Resulting phylogenies were uploaded into the ITOL program (Letunic and Bork, 2011) for graphical display of family and twinID annotations, and into Geneious (v 6.1.8) to retrieve patristic distances (branch lengths). Mean patristic distances were compared across comparison categories using bootstrapped Wilcoxon rank sum tests as described above.

All software was run parallelized on a high performance computing cluster at the Max Planck Institute for Developmental Biology via Snakemake (Köster and Rahmann, 2012).

### Synteny analyses

First, a de-novo assembly of post-captured metagenomes was performed for each sample by using the metacompass software (Cepeda *et al.*, 2017). Next, a set of genes representing different genomic regions was selected randomly (using a custom script) from each of the reference genomes of *B. longum, B. adolescentis, B. animalis* and *B.bifidum.* Each gene was used as a query for a BLASTn (Altschul *et al.*, 1990) search against the assembled metagenomes, with minimal identity of 97% and minimal coverage of 90%. For each of the blast hits in the assembled metagenomes, the gene and flanking sequences (3.5kb upstream and downstream to the blast hit) were retrieved and used for synteny comparison. Further analysis was only carried out for regions found in > 15 samples. If for a given species the final number of suitable regions was lower than 4, the process iterated with a new set of genes, keeping a minimal gap of 5 genes between each two selected genes, to avoid overlap between regions.

Pairwise synteny comparison between each two DNA sequences was done using the DECIPHER R package (Wright, 2016). Breaks in the synteny were defined as non-homologous regions longer than 15 base pairs. To compare the synteny between different types of pairwise comparisons (i.e. intraindividual, twins, non-related individuals) we defined the synteny score (equation 1):

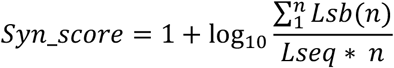

Where ***n*** is the number of synteny blocks identified in each pairwise comparison, ***Lseq*** is the length of the shorter sequence in each pair of compared sequences and ***Lsb(n)*** represents the length of the ***n***^th^ synteny block.

## DECLERATIONS

### Ethics approval and consent to participate

All work involving the use of these previously collected samples from human subjects was approved by the Cornell University IRB (Protocol ID 1108002388).

### Consent for publication

Not applicable.

### Availability of data and materials

Sequences of pre and post captured metagenomes (the only new data generated for this manuscript) are available on the European Nucleotide Archive accession number PRJEB38000.

### Funding

This work was supported by the Max Planck Society and NIH grant# 2R01DK093595-05A1

### Authors’ contributions

VS and RL conceived the project. VS and NY ran the bioinformatics and statistical analyses. HE conceived and executed Synteny analyses. VS, RL, HE wrote the paper. RL and TS supervised the project.

## Acknowledgements

We thank the team at NimbleGene and Roche Sequencing Solutions, Nicole Leahy, Christine Kuch, and others, for their help in developing the genome capture array. TwinsUK is funded by the Wellcome Trust, Medical Research Council, European Union, Chronic Disease Research Foundation (CDRF), Zoe Global Ltd and the National Institute for Health Research (NIHR)-funded BioResource, Clinical Research Facility and Biomedical Research Centre based at Guy’s and St Thomas’ NHS Foundation Trust in partnership with King’s College London.

## ABBREVIATIONS

MLSA: Multilocus sequence alignment
SV: Sequence Variant (a.k.a ASV or 100% OTU)
BCD: Bray-Curtis Dissimilarity
MIDAS: Metagenomic Intra-Species Diversity Analysis System

